## Supplementary material for "Close-kin mark-recapture methods to estimate demographic parameters of mosquitoes": S1 Text

### Close-kin mark-recapture methods to estimate demographic parameters of mosquitoes
### S1 Text: Supplemental model equations


Yogita Sharma[1+], Jared B. Bennett[2+], Gordana Rašić[3], and John M. Marshall[1,4*]

[1]Divisions of Biostatistics and Epidemiology, School of Public Health, University of California, Berkeley, CA, USA
[2]Biophysics Graduate Group, Division of Biological Sciences, College of Letters and Science, University of California, Berkeley, CA, USA
[3]Mosquito Genomics, QIMR Berghofer Medical Research Institute, Brisbane, Australia
[4]Innovative Genomics Institute, University of California, Berkeley, CA, USA
[+]Equal contribution



August 2022


#### 1 Mosquito population dynamics:

In the manuscript, we describe a discrete-time version of the lumped age-class model (Gurney *et al.*, 1983) applied to mosquitoes (Hancock & Godfray, 2007) as a basis for our population simulation and close-kin mark-recapture (CKMR) analysis. This model considers discrete life history stages - egg (E), larva (L), pupa (P) and adult (A) - with sub-adult stages having defined durations - $T_E$, $T_L$ and $T_P$ for eggs, larvae and pupae, respectively. Additional terms are defined in the manuscript (§2.1). Here, we include equations and derivations that, for brevity, were not included in the manuscript.

##### 1.1 Close-kin mark-recapture analysis:

Of relevance to the close-kin mark-recapture (CKMR) analysis, here we show how: i) juvenile mortality rates are derived from the daily mosquito population growth rate, $r_M$, and ii) the larval mortality rate, $\mu_L$, is be chosen to produce a population at equilibrium. The derivations in this section will be based on the generational population growth rate, $R_M$. This is related to the daily population growth rate, $r_M$, according to:

$$R_M = (r_M)^G.$$

Here, $G$ represents the mosquito generation time, which is equal to the sum of the juvenile life stage durations and the life expectancy of the adult stage:

$$G = T_E + T_L + T_P + (1/\mu_A).$$

The generational population growth rate is equal to the daily adult survival rate multiplied by half the female egg production rate (because half the adults are female) multiplied by the proportion of eggs that survive through all of the juvenile life stages in the absence of density-dependence, multiplied by the adult life expectancy:

$$R_M = \frac{\beta}{2} \times (1-\mu_E)^{T_E} \times (1-\mu_L)^{T_L} \times (1-\mu_P)^{T_P} \times \frac{(1-\mu_A)}{\mu_A}.$$

Juvenile mortality rates are difficult to measure in the wild, and so in considering the case of density-independent population growth, we treat the egg, larval and pupal daily mortality rates as all being equal to the juvenile mortality rate, $\mu_J$. The equation for $R_M$ therefore simplifies to:

$$R_M = \frac{\beta \times (1-\mu_J)^{(T_E+T_L+T_P)} \times (1-\mu_A)}{2\mu_A}.$$

This can then be rearranged to to obtain the density-independent juvenile mortality rate, $\mu_J$, in terms of the generational population growth rate, $R_M$:

$$\mu_J = 1 - \left(\frac{2R_M\mu_A}{\beta(1-\mu_A)}\right)^{(1/(T_E+T_L+T_P))}.$$

As with many other mosquito population dynamic models, we assume that population regulation occurs at the larval stage, moderated by a density-dependent larval mortality rate. While we hold the rates of egg and pupal mortality equal to the density-independent juvenile mortality rate, we calculate the density-dependent larval mortality rate, $\mu_L$, that produces a population at equilibrium by setting $R_M$ equal to one, i.e.:

$$R_M = 1 = \frac{\beta \times (1-\mu_J)^{(T_E+T_P)} \times (1-\mu_L)^{(T_L)} \times (1-\mu_A)}{2\mu_A}.$$

This can then be rearranged to to obtain the density-dependent larval mortality rate:

$$\mu_L = 1 - \left(\frac{2\mu_A}{\beta(1-\mu_A)(1-\mu_J)^{(T_E+T_P)}}\right)^{(1/T_L)}.$$

For *Aedes aegypti*, we obtain a daily population growth rate, $r_M$, of 1.175 per day (Simoy *et al.*, 2015). From the equation for $\mu_J$, this results in an egg and pupal mortality rate of 0.175 per day, and from the above equation, a larval mortality rate of 0.554 per day results in a population at equilibrium.

#### 1.2 Individual-based simulation model:

For the CKMR analysis, we assume a population at equilibrium, and so calculate a larval mortality rate, $\mu_L$, that produces this equilibrium. The individual-based simulation model, on the other hand, is stochastic, and so the density-dependent component of the larval mortality rate must be dynamic and respond to the current state of the population. Following Deredec *et al.* (2011), we model density-dependence at the larval stage by reducing the density-independent daily larval survival probability by a fraction, $\alpha/(\alpha + N_L)$, where $N_L$ represents the larval population size, and $\alpha$ is a parameter influencing the strength of density-dependence. Considering $\mu_J$ as the density-independent juvenile (and larval) mortality rate, the density-dependent daily larval survival probability is therefore:

$$(1-\mu_L) = (1-\mu_J) \times \frac{\alpha}{\alpha + N_L}.$$

This form of density dependence is a discrete-time version of the logistic growth model used by Berverton & Holt (1957). The value of $\alpha$ is chosen to produce the desired equilibrium adult population size, $N_A^*$, and can be shown to be equal to:

$$\alpha = \frac{\beta \times (1-\mu_J) \times N_A^*}{2 \times (R_M - 1)} \times \frac{1 - ((1-\mu_J)/R_M)}{1 - ((1-\mu_J)/R_M)^{1/T_L}}.$$

### 2 Kinship probabilities:

Here, we include the kinship probabilities for parent-offspring and sibling pairs that, for brevity, were not included in the manuscript (§2.2).

#### 2.1 Mother-offspring:

First, we adapt the mother-offspring kinship probability for adult offspring to obtain $P_{MOA}(t_2|t_1)$, the probability that, given an adult female sampled on day $t_1$, an adult sampled on day $t_2$ is her offspring:

$$P_{MOA}(t_2|t_1) = \frac{\mathbb{E}\left[\text{Adult offspring at time } t_2 \text{ from an adult female sampled at time } t_1\right]}{\mathbb{E}\left[\text{Adult offspring at time } t_2 \text{ from all adult females}\right]} = \frac{E_{MOA}(t_2|t_1)}{E_A}.$$

Here, $E_{MOA}(t_2|t_1)$ represents the expected number of surviving adult offspring on day $t_2$ from an adult female sampled on day $t_1$, and is given by:

$$E_{MOA}(t_2|t_1) = \sum_{y_2=t_2-T_E-T_L-T_P-(T_A-1)}^{t_2-T_E-T_L-T_P} (1-\mu_A)^{(t_1-y_2)} \times \begin{pmatrix} \mathbb{I}[(t_1 - T_A) < y_2 \leq t_1] \times \beta \times (1-\mu_E)^{T_E} \\ \times (1-\mu_L)^{T_L} \times (1-\mu_P)^{T_P} \\ \times (1-\mu_A)^{(t_2-y_2-T_E-T_L-T_P)} \end{pmatrix}.$$

Here, the day of egg-laying, $y_2$, is summed over days $(t_2 - T_E - T_L - T_P - (T_A - 1))$ through $(t_2 - T_E - T_L - T_P)$, for consistency with the adult offspring being present on the day of sampling. The terms within the summation are the same as for the mother-larval offspring case, with the exception that daily egg production is multiplied by the proportion of eggs that survive the egg, larva, pupa and adult stages from the day they were laid up to the day of sampling, $t_2$.

Next, we adapt the mother-offspring kinship probability for pupal offspring to obtain $P_{MOP}(t_2|t_1)$, the probability that, given an adult female sampled on day $t_1$, a pupa sampled on day $t_2$ is her offspring:

$$P_{MOP}(t_2|t_1) = \frac{\mathbb{E}\left[\text{Pupal offspring at time } t_2 \text{ from an adult female sampled at time } t_1\right]}{\mathbb{E}\left[\text{Pupal offspring at time } t_2 \text{ from all adult females}\right]} = \frac{E_{MOP}(t_2|t_1)}{E_P}.$$

Here, $E_{MOP}(t_2|t_1)$ represents the expected number of surviving pupal offspring on day $t_2$ from an adult female sampled on day $t_1$, and $E_P$ represents the expected number of surviving pupal offspring from all adult females at times consistent with the time of pupal sampling. Assuming a population at equilibrium, $E_P$ is independent of time and is given by:

$$E_P = \sum_{y_2=0-T_E-T_L-(T_P-1)}^{0-T_E-T_L} N \times \beta \times (1-\mu_E)^{T_E} \times (1-\mu_L)^{T_L} \times (1-\mu_P)^{(0-y_2-T_E-T_L)}.$$

Here, considering day 0 as the reference day, the day of egg-laying, $y_2$, is summed over days $(t_2 - T_E - T_L - (T_P - 1))$ through $(t_2 - T_E - T_L)$, for consistency with the pupal offspring being present on the day of sampling. This equation therefore represents the expected number of offspring laid by all adult females in the population that survive the egg, larva and pupa stages up to the time of sampling (day 0). $E_{MOP}(t_2|t_1)$ is then given by:

$$E_{MOP}(t_2|t_1) = \sum_{y_2=t_2-T_E-T_L-(T_P-1)}^{t_2-T_E-T_L} (1-\mu_A)^{(t_1-y_2)} \times \begin{pmatrix} \mathbb{I}[(t_1-T_A) < y_2 \leq t_1] \times \beta \times (1-\mu_E)^{T_E} \\ \times (1-\mu_L)^{T_L} \times (1-\mu_P)^{(t_2-y_2-T_E-T_L)} \end{pmatrix}.$$

Here, the day of egg-laying, $y_2$, is summed over days $(t_2 - T_E - T_L - (T_P - 1))$ through $(t_2 - T_E - T_L)$, for consistency with the pupal offspring being present on the day of sampling. The terms within the summation are then the same as for the mother-larval offspring case, with the exception that daily egg production is multiplied by the proportion of eggs that survive the egg, larva and pupa stages from the day they were laid up to the day of sampling, $t_2$.

#### 2.2 Father-offspring:

Next, we adapt the father-offspring kinship probability for larval offspring to obtain $P_{FOL}(t_2|t_1)$, the probability that, given an adult male sampled on day $t_1$, a larva sampled on day $t_2$ is his offspring:

$$P_{FOL}(t_2|t_1) = \frac{\mathbb{E}\left[\text{Larval offspring at time } t_2 \text{ from an adult male sampled at time } t_1\right]}{\mathbb{E}\left[\text{Larval offspring at time } t_2 \text{ from all adult females}\right]} = \frac{E_{FOL}(t_2|t_1)}{E_L}.$$

Here, $E_{FOL}(t_2|t_1)$ represents the expected number of surviving larval offspring on day $t_2$ of an adult male sampled on day $t_1$. Each adult male mates on average once; but the day of this mating event, $t_i$, is unknown and so, in calculating $E_{FOL}(t_2|t_1)$, we treat this as a latent variable and take an expectation over all possible values it can take:

$$E_{FOL}(t_2|t_1) = \sum_{t_i=t_1-(T_A-1)}^{t_1} p_A(t_1-t_i) \times E_{FOL}(t_2|t_1,t_i).$$

Here, the expectation over the day of mating, $t_i$, is taken over days $(t_1 - (T_A - 1))$ through $t_1$, for consistency with the day of adult male sampling. The term $E_{FOL}(t_2|t_1, t_i)$ represents the expected number of surviving larval offspring on day $t_2$ of an adult male sampled on day $t_1$, conditional upon the day of mating being $t_i$, and is given by:

$$E_{FOL}(t_2|t_1,t_i) = \sum_{y_2=t_i}^{t_i+(T_A-1)} (1-\mu_A)^{(y_2-t_i)} \times \begin{pmatrix} \mathbb{I}[(y_2+T_E) < t_2 \leq (y_2+T_E+T_L)] \times \\ \beta \times (1-\mu_E)^{T_E} \times (1-\mu_L)^{(t_2-y_2-T_E)} \end{pmatrix}.$$

Here, the day of egg-laying, $y_2$, is summed over days $t_i$ through $(t_i + (T_A - 1))$, for consistency with the mother's potential lifespan. The terms within the summation are then the same as for the father-adult offspring case, with the exception that daily egg production is multiplied by the proportion of eggs that survive the egg and larva stages from the day they were laid up to the day of sampling, $t_2$, and the indicator function limits consideration to cases where the day of larval sampling lies within the larval offspring's possible lifetime - i.e. between days $(y_2 + T_E)$ and $(y_2 + T_E + T_L)$.

Next, we adapt the father-offspring kinship probability for pupal offspring to obtain $P_{FOP}(t_2|t_1)$, the probability that, given an adult male sampled on day $t_1$, a pupa sampled on day $t_2$ is his offspring:

$$P_{FOP}(t_2|t_1) = \frac{\mathbb{E}\left[\text{Pupal offspring at time } t_2 \text{ from an adult male sampled at time } t_1\right]}{\mathbb{E}\left[\text{Pupal offspring at time } t_2 \text{ from all adult females}\right]} = \frac{E_{FOP}(t_2|t_1)}{E_P}.$$

Here, $E_{FOP}(t_2|t_1)$ represents the expected number of surviving pupal offspring on day $t_2$ of an adult male sampled on day $t_1$. Each adult male mates on average once; but the day of this mating event, $t_i$, is unknown and so, in calculating $E_{FOP}(t_2|t_1)$, we treat this as a latent variable and take an expectation over all possible values it can take:

$$E_{FOP}(t_2|t_1) = \sum_{t_i = t_1 - (T_A - 1)}^{t_1} p_A(t_1 - t_i) \times E_{FOP}(t_2|t_1, t_i).$$

Here, the expectation over the day of mating, $t_i$, is taken over days $(t_1 - (T_A - 1))$ through $t_1$, for consistency with the day of adult male sampling. The term $E_{FOP}(t_2|t_1, t_i)$ represents the expected number of surviving pupal offspring on day $t_2$ of an adult male sampled on day $t_1$, conditional upon the day of mating being $t_i$, and is given by:

$$E_{FOP}(t_2|t_1, t_i) = \sum_{y_2 = t_i}^{t_i + (T_A - 1)} (1 - \mu_A)^{(y_2 - t_i)} \times \begin{pmatrix} \mathbb{I}[(y_2 + T_E + T_L) \leq t_2 < (y_2 + T_E + T_L + T_P)] \times \beta \\ \times (1 - \mu_E)^{T_E} \times (1 - \mu_L)^{T_L} \times (1 - \mu_P)^{(t_2 - y_2 - T_E - T_L)} \end{pmatrix}.$$

Here, the day of egg-laying, $y_2$, is summed over days $t_i$ through $(t_i + (T_A - 1))$, for consistency with the mother's potential lifespan. The terms within the summation are then the same as for the father-adult offspring case, with the exception that daily egg production is multiplied by the proportion of eggs that survive the egg, larva and pupa stages from the day they were laid up to the day of sampling, $t_2$, and the indicator function limits consideration to cases where the day of pupal sampling lies within the adult offspring's possible lifetime - i.e. between days $(y_2 + T_E + T_L)$ and $(y_2 + T_E + T_L + T_P)$.

#### 2.3 Full-siblings:

Next, we consider the adult-adult full-sibling kinship probability, $P_{FSAA}(t_2|t_1)$, which represents the probability that, given an adult sampled on day $t_1$, an adult sampled on day $t_2$ is their full-sibling:

$$P_{FSAA}(t_2|t_1) = \frac{\mathbb{E}\left[\text{Adults at time } t_2 \text{ that are full-siblings of an adult sampled at time } t_1\right]}{\mathbb{E}\left[\text{Adult offspring at time } t_2 \text{ from all adult females}\right]} = \frac{E_{FSAA}(t_2|t_1)}{E_A}.$$

Here, $E_{FSAA}(t_2|t_1)$ represents the expected number of surviving adults on day $t_2$ that are full-siblings of an adult sampled on day $t_1$. For convenience, let us refer to the adult sampled on day $t_1$ as individual 1.

To calculate $E_{FSAA}(t_1, t_2)$, there are two unknown event times that we treat as latent variables and take an expectation over - i) the day that egg 1 is laid, $y_1$, and ii) the day that individual 1's mother emerges as an adult, $t_i$:

$$E_{FSAA}(t_2|t_1) = \sum_{y_1=t_1-T_E-T_L-T_P-(T_A-1)}^{t_1-T_E-T_L-T_P} p_A(t_1-y_1-T_E-T_L-T_P) \times \sum_{t_i=y_1-(T_A-1)}^{y_1} p_A(y_1-t_i) \times E_{FSAA}(t_2|t_1, y_1, t_i).$$

This is the same equation as for the larva-larva case with two exceptions: i) the expectation over the day that individual 1 is laid is taken over days $(t_1 - T_E - T_L - T_P - (T_A - 1))$ through $(t_1 - T_E - T_L - T_P)$ to account for the additional time elapsed between the larva and adult life stages, and ii) the probability that egg 1 is laid on day $(t_1 - y_1 - T_E - T_L - T_P)$, $p_A(t_1 - y_1 - T_E - T_L - T_P)$, reflects the adult age probability distribution as this is the relevant life stage. The term $E_{FSAA}(t_2|t_1, y_1, t_i)$ represents the expected number of surviving adults on day $t_2$ that are full-siblings of adult 1, conditional upon egg 1 being laid on day $y_1$, and their mother emerging as an adult on day $t_i$. This is given by:

$$E_{FSAA}(t_2|t_1, y_1, t_i) = \sum_{y_2=t_i}^{t_i+(T_A-1)} (1-\mu_A)^{(y_2-t_i)} \times \begin{pmatrix} \mathbb{I}[(t_2 - T_E - T_L - T_P - T_A) < y_2 \leq (t_2 - T_E - T_L - T_P)] \\ \times \beta \times (1 - \mu_E)^{T_E} \times (1 - \mu_L)^{T_L} \\ \times (1 - \mu_P)^{T_P} \times (1 - \mu_A)^{(t_2-y_2-T_E-T_L-T_P)} \end{pmatrix}.$$

This is the same equation as for the larva-larva case with two exceptions: i) daily egg production is multiplied by the proportion of eggs that survive the egg, larva, pupa and adult stages up to the day of sampling to reflect the fact that adults rather than larvae are being sampled, and ii) the indicator function limits consideration to cases where the day of sibling egg-laying, $y_2$, is between days $(t_2 - T_E - T_L - T_P - T_A)$ and $(t_2 - T_E - T_L - T_P)$, again accounting for the additional time elapsed between the larva and adult life stages.

Next, we consider the larva-adult full-sibling kinship probability, $P_{FSLA}(t_2|t_1)$, which represents the probability that, given a larva sampled on day $t_1$, an adult sampled on day $t_2$ is their full-sibling:

$$P_{FSLA}(t_2|t_1) = \frac{\mathbb{E}\left[\text{Adults at time } t_2 \text{ that are full-siblings of a larva sampled at time } t_1\right]}{\mathbb{E}\left[\text{Adult offspring at time } t_2 \text{ from all adult females}\right]} = \frac{E_{FSLA}(t_2|t_1)}{E_A}.$$

Here, $E_{FSLA}(t_2|t_1)$ represents the expected number of surviving adults on day $t_2$ that are full-siblings of a larva sampled on day $t_1$. For convenience, let us refer to the larva sampled on day $t_1$ as individual 1. To calculate $E_{FSLA}(t_2|t_1)$, there are two unknown event times that we treat as latent variables and take an expectation over - i) the day that egg 1 is laid, $y_1$, and ii) the day that individual 1's mother emerges as an adult, $t_i$:

$$E_{FSLA}(t_2|t_1) = \sum_{y_1=t_1-T_E-(T_L-1)}^{t_1-T_E} p_L(t_1 - y_1 - T_E) \times \sum_{t_i=y_1-(T_A-1)}^{y_1} p_A(y_1 - t_i) \times E_{FSLA}(t_2|t_1, y_1, t_i).$$

This is the same equation as for the larva-larva case with the only exception being the term $P_{FSLA}(t_2|t_1, y_1, t_i)$, which represents the expected number of surviving adults on day $t_2$ that are full-siblings of larva 1, conditional upon egg 1 being laid on day $y_1$, and their mother emerging as an adult on day $t_i$. That term is the same as for the adult-adult case, i.e.:

$$E_{FSLA}(t_2|t_1, y_1, t_i) = E_{FSAA}(t_2|t_1, y_1, t_i).$$

Next, we consider the adult-larva full-sibling kinship probability, $P_{FSAL}(t_2|t_1)$, which represents the probability that, given an adult sampled on day $t_1$, a larva sampled on day $t_2$ is their full-sibling:

$$P_{FSAL}(t_2|t_1) = \frac{\mathbb{E}\left[\text{Larvae at time } t_2 \text{ that are full-siblings of an adult sampled at time } t_1\right]}{\mathbb{E}\left[\text{Larval offspring at time } t_2 \text{ from all adult females}\right]} = \frac{E_{FSAL}(t_2|t_1)}{E_L}.$$

Here, $E_{FSAL}(t_2|t_1)$ represents the expected number of surviving larvae on day $t_2$ that are full-siblings of an adult sampled on day $t_1$. For convenience, let us refer to the adult sampled on day $t_1$ as individual 1. To calculate $E_{FSAL}(t_2|t_1)$, there are two unknown event times that we treat as latent variables and take an expectation over - i) the day that egg 1 is laid, $y_1$, and ii) the day that individual 1's mother emerges as an adult, $t_i$:

$$E_{FSAL}(t_2|t_1) = \sum_{y_1=t_1-T_E-T_L-T_P-(T_A-1)}^{t_1-T_E-T_L-T_P} p_A(t_1-y_1-T_E-T_L-T_P) \times \sum_{t_i=y_1-(T_A-1)}^{y_1} p_A(y_1-t_i) \times E_{FSAL}(t_2|t_1, y_1, t_i).$$

This is the same equation as for the adult-adult case with the only exception being the term $E_{FSAL}(t_2|t_1, y_1, t_i)$, which represents the expected number of surviving larvae on day $t_2$ that are full-siblings of adult 1, conditional upon egg 1 being laid on day $y_1$, and their mother emerging as an adult on day $t_i$. That term is the same as for the larva-larva case, i.e.:

$$E_{FSAL}(t_2|t_1, y_1, t_i) = E_{FSLL}(t_2|t_1, y_1, t_i).$$

For pupa-pupa pairs, the full-sibling kinship probability is denoted by $P_{FSPP}(t_2|t_1)$ and represents the probability that, given a pupa sampled on day $t_1$, a pupa sampled on day $t_2$ is their full-sibling. This can be expressed as:

$$P_{FSPP}(t_2|t_1) = \frac{\mathbb{E}\left[\text{Pupae at time } t_2 \text{ that are full-siblings of a pupa sampled at time } t_1\right]}{\mathbb{E}\left[\text{Pupal offspring at time } t_2 \text{ from all adult females}\right]} = \frac{E_{FSPP}(t_2|t_1)}{E_P}.$$

Here, $E_{FSPP}(t_2|t_1)$ represents the expected number of surviving pupae on day $t_2$ that are full-siblings of a pupa sampled on day $t_1$. For convenience, let us refer to the pupa sampled on day $t_1$ as individual 1. To calculate $E_{FSPP}(t_2|t_1)$, there are two unknown event times that we treat as latent variables and take an expectation over - i) the day that egg 1 is laid, $y_1$, and ii) the day that individual 1's mother emerges as an adult, $t_i$:

$$E_{FSPP}(t_2|t_1) = \sum_{y_1=t_1-T_E-T_L-(T_P-1)}^{t_1-T_E-T_L} p_P(t_1 - y_1 - T_E - T_L) \times \sum_{t_i=y_1-(T_A-1)}^{y_1} p_A(y_1 - t_i) \times E_{FSPP}(t_2|t_1, y_1, t_i).$$

This is the same equation as for the larva-larva case with two exceptions: i) the expectation over the day that individual 1 is laid is taken over days $(t_1 - T_E - T_L - (T_P - 1))$ through $(t_1 - T_E - T_L)$ to account for the additional time elapsed between the larva and pupa life stages, and ii) the probability that egg 1 is laid on day $(t_1 - y_1 - T_E - T_L)$, $p_P(t_1 - y_1 - T_E - T_L)$, reflects the pupa age probability distribution due to this being the relevant life stage. In general, $p_P(t)$ represents the probability that a given pupa in the population has age $t$ which, following from the daily pupal survival probability, $(1 - \mu_P)$, is given by:

$$p_P(t) = (1 - \mu_P)^t \Big/ \sum_{t_j=0}^{T_P-1} (1 - \mu_P)^{t_j}.$$

The term $E_{FSPP}(t_2|t_1, y_1, t_i)$ represents the expected number of surviving pupae on day $t_2$ that are full-siblings of pupa 1, conditional upon egg 1 being laid on day $y_1$, and their mother emerging as an adult on day $t_i$. This is given by:

$$E_{FSPP}(t_2|t_1, y_1, t_i) = \sum_{y_2=t_i}^{t_i+(T_A-1)} (1 - \mu_A)^{(y_2-t_i)} \times \begin{pmatrix} \mathbb{I}[(t_2 - T_E - T_L - T_P) < y_2 \leq (t_2 - T_E - T_L)] \times \beta \\ \times (1 - \mu_E)^{T_E} \times (1 - \mu_L)^{T_L} \times (1 - \mu_P)^{(t_2-y_2-T_E-T_L)} \end{pmatrix}.$$

This is the same equation as for the larva-larva case with two exceptions: i) daily egg production is multiplied by the proportion of eggs that survive the egg, larva and pupa stages up to the day of sampling to reflect the fact that pupae rather than larvae are being sampled, and ii) the indicator function limits consideration to cases where the day of sibling egg-laying, $y_2$, is between days $(t_2 - T_E - T_L - T_P)$ and $(t_2 - T_E - T_L)$, again accounting for the additional time elapsed between the larva and pupa life stages.

Next, we consider the pupa-larva full-sibling kinship probability, $P_{FSPL}(t_2|t_1)$, which represents the probability that, given a pupa sampled on day $t_1$, a larva sampled on day $t_2$ is their full-sibling:

$$P_{FSPL}(t_2|t_1) = \frac{\mathbb{E}\left[\text{Larvae at time } t_2 \text{ that are full-siblings of a pupa sampled at time } t_1\right]}{\mathbb{E}\left[\text{Larval offspring at time } t_2 \text{ from all adult females}\right]} = \frac{E_{FSPL}(t_2|t_1)}{E_L}.$$

Here, $E_{FSPL}(t_2|t_1)$ represents the expected number of surviving larvae on day $t_2$ that are full-siblings of a pupa sampled on day $t_1$. For convenience, let us refer to the pupa sampled on day $t_1$ as individual 1. To calculate $E_{FSPL}(t_2|t_1)$, there are two unknown event times that we treat as latent variables and take an expectation over - i) the day that egg 1 is laid, $y_1$, and ii) the day that individual 1's mother emerges as an adult, $t_i$:

$$E_{FSPL}(t_2|t_1) = \sum_{y_1=t_1-T_E-T_L-(T_P-1)}^{t_1-T_E-T_L} p_P(t_1 - y_1 - T_E - T_L) \times \sum_{t_i=y_1-(T_A-1)}^{y_1} p_A(y_1 - t_i) \times E_{FSPL}(t_2|t_1, y_1, t_i).$$

This is the same equation as for the pupa-pupa case with the only exception being the term $E_{FSPL}(t_2|t_1, y_1, t_i)$, which represents the expected number of surviving larvae on day $t_2$ that are full-siblings of pupa 1, conditional upon egg 1 being laid on day $y_1$, and their mother emerging as an adult on day $t_i$. That term is the same as for the larva-larva case, i.e.:

$$E_{FSPL}(t_2|t_1, y_1, t_i) = E_{FSLL}(t_2|t_1, y_1, t_i).$$

Next, we consider the larva-pupa full-sibling kinship probability, $P_{FSLP}(t_2|t_1)$, which represents the probability that, given a larva sampled on day $t_1$, a pupa sampled on day $t_2$ is their full-sibling:

$$P_{FSLP}(t_2|t_1) = \frac{\mathbb{E}\left[\text{Pupae at time } t_2 \text{ that are full-siblings of a larva sampled at time } t_1\right]}{\mathbb{E}\left[\text{Pupal offspring at time } t_2 \text{ from all adult females}\right]} = \frac{E_{FSLP}(t_2|t_1)}{E_P}.$$

Here, $E_{FSLP}(t_2|t_1)$ represents the expected number of surviving pupae on day $t_2$ that are full-siblings of a larva sampled on day $t_1$. For convenience, let us refer to the larva sampled on day $t_1$ as individual 1. To calculate $E_{FSLP}(t_2|t_1)$, there are two unknown event times that we treat as latent variables and take an expectation over - i) the day that egg 1 is laid, $y_1$, and ii) the day that individual 1's mother emerges as an adult, $t_i$:

$$E_{FSLP}(t_2|t_1) = \sum_{y_1=t_1-T_E-(T_L-1)}^{t_1-T_E} p_L(t_1 - y_1 - T_E) \times \sum_{t_i=y_1-(T_A-1)}^{y_1} p_A(y_1 - t_i) \times E_{FSLP}(t_2|t_1, y_1, t_i).$$

This is the same equation as for the larva-larva case with the only exception being the term $E_{FSLP}(t_2|t_1, y_1, t_i)$, which represents the expected number of surviving pupae on day $t_2$ that are full-siblings of larva 1, conditional upon egg 1 being laid on day $y_1$, and their mother emerging as an adult on day $t_i$. That term is the same as for the pupa-pupa case, i.e.:

$$E_{FSLP}(t_2|t_1, y_1, t_i) = E_{FSPP}(t_2|t_1, y_1, t_i).$$

Next, we consider the pupa-adult full-sibling kinship probability, $P_{FSPA}(t_2|t_1)$, which represents the probability that, given a pupa sampled on day $t_1$, an adult sampled on day $t_2$ is their full-sibling:

$$P_{FSPA}(t_2|t_1) = \frac{\mathbb{E}\left[\text{Adults at time } t_2 \text{ that are full-siblings of a pupa sampled at time } t_1\right]}{\mathbb{E}\left[\text{Adult offspring at time } t_2 \text{ from all adult females}\right]} = \frac{E_{FSPA}(t_2|t_1)}{E_A}.$$

Here, $E_{FSPA}(t_2|t_1)$ represents the expected number of surviving adults on day $t_2$ that are full-siblings of a pupa sampled on day $t_1$. For convenience, let us refer to the pupa sampled on day $t_1$ as individual 1. To calculate $E_{FSPA}(t_2|t_1)$, there are two unknown event times that we treat as latent variables and take an expectation over - i) the day that egg 1 is laid, $y_1$, and ii) the day that individual 1's mother emerges as an adult, $t_i$:

$$E_{FSPA}(t_2|t_1) = \sum_{y_1=t_1-T_E-T_L-(T_P-1)}^{t_1-T_E-T_L} p_P(t_1 - y_1 - T_E - T_L) \times \sum_{t_i=y_1-(T_A-1)}^{y_1} p_A(y_1 - t_i) \times E_{FSPA}(t_2|t_1, y_1, t_i).$$

This is the same equation as for the pupa-pupa case with the only exception being the term $E_{FSPA}(t_2|t_1, y_1, t_i)$, which represents the expected number of surviving adults on day $t_2$ that are full-siblings of pupa 1, conditional upon egg 1 being laid on day $y_1$, and their mother emerging as an adult on day $t_i$. That term is the same as for the adult-adult case, i.e.:

$$E_{FSPA}(t_2|t_1, y_1, t_i) = E_{FSAA}(t_2|t_1, y_1, t_i).$$

Next, we consider the adult-pupa full-sibling kinship probability, $P_{FSAP}(t_2|t_1)$, which represents the probability that, given an adult sampled on day $t_1$, a pupa sampled on day $t_2$ is their full-sibling:

$$P_{FSAP}(t_2|t_1) = \frac{\mathbb{E}\left[\text{Pupae at time } t_2 \text{ that are full-siblings of an adult sampled at time } t_1\right]}{\mathbb{E}\left[\text{Pupal offspring at time } t_2 \text{ from all adult females}\right]} = \frac{E_{FSAP}(t_2|t_1)}{E_P}.$$

Here, $E_{FSAP}(t_2|t_1)$ represents the expected number of surviving pupae on day $t_2$ that are full-siblings of an adult sampled on day $t_1$. For convenience, let us refer to the adult sampled on day $t_1$ as individual 1. To calculate $E_{FSAP}(t_2|t_1)$, there are two unknown event times that we treat as latent variables and take an expectation over - i) the day that egg 1 is laid, $y_1$, and ii) the day that individual 1's mother emerges as an adult, $t_i$:

$$E_{FSAP}(t_2|t_1) = \sum_{y_1=t_1-T_E-T_L-T_P-(T_A-1)}^{t_1-T_E-T_L-T_P} p_A(t_1-y_1-T_E-T_L-T_P)\times \sum_{t_i=y_1-(T_A-1)}^{y_1} p_A(y_1-t_i)\times E_{FSAP}(t_2|t_1,y_1,t_i).$$

This is the same equation as for the adult-adult case with the only exception being the term $E_{FSAP}(t_2|t_1,y_1,t_i)$, which represents the expected number of surviving pupae on day $t_2$ that are full-siblings of adult 1, conditional upon egg 1 being laid on day $y_1$, and their mother emerging as an adult on day $t_i$. That term is the same as for the pupa-pupa case, i.e.:

$$E_{FSAP}(t_2|t_1,y_1,t_i) = E_{FSPP}(t_2|t_1,y_1,t_i).$$

#### 2.4 Half-siblings:

Next, we consider the adult-adult half-sibling kinship probability, $P_{HSAA}(t_2|t_1)$, which represents the probability that, given an adult sampled on day $t_1$, an adult sampled on day $t_2$ is their half-sibling:

$$P_{HSAA}(t_2|t_1) = \frac{\mathbb{E}\left[\text{Adults at time } t_2 \text{ that are half-siblings of an adult sampled at time } t_1\right]}{\mathbb{E}\left[\text{Adult offspring at time } t_2 \text{ from all adult females}\right]} = \frac{E_{HSAA}(t_2|t_1)}{E_A}.$$

Here, $E_{HSAA}(t_2|t_1)$ represents the expected number of surviving adults on day $t_2$ that are half-siblings of an adult sampled on day $t_1$. For convenience, let us refer to the adult sampled on day $t_1$ as individual 1. To calculate $E_{HSAA}(t_2|t_1)$, there are three unknown event times that we treat as latent variables and take an expectation over - i) the day that egg 1 is laid, $y_1$, ii) the day of the mating event between individual 1's mother and father, $t_i$, and iii) the day that individual 1's father emerges as an adult, $t_j$:

$$\begin{aligned} E_{HSAA}(t_2|t_1) = &\sum_{y_1=t_1-T_E-T_L-T_P-(T_A-1)}^{t_1-T_E-T_L-T_P} p_A(t_1-y_1-T_E-T_L-T_P)\times \sum_{t_i=y_1-(T_A-1)}^{y_1} p_A(y_1-t_i) \\ &\times \sum_{t_j=t_i-(T_A-1)}^{t_i} p_A(t_i-t_j)\times E_{HSAA}(t_2|t_1,y_1,t_i,t_j). \end{aligned}$$

This is the same equation as for the larva-larva case with two exceptions: i) the expectation over the day that individual 1 is laid is taken over days $(t_1-T_E-T_L-T_P-(T_A-1))$ through $(t_1-T_E-T_L-T_P)$ to account for the additional time elapsed between the larva and adult life stages, and ii) the probability that egg 1 is laid on day $(t_1-y_1-T_E-T_L-T_P)$, $p_A(t_1-y_1-T_E-T_L-T_P)$, reflects the adult age probability distribution due to this being the relevant life stage. The term $E_{HSAA}(t_2|t_1, y_1, t_i, t_j)$ represents the expected number of surviving adults on day $t_2$ that are half-siblings of adult 1, conditional upon egg 1 being laid on day $y_1$, their mother and father mating on day $t_i$, and their father emerging as an adult on day $t_j$. This is given by:

$$E_{HSAA}(t_2|t_1, y_1, t_i, t_j) = \sum_{t_k=t_j}^{t_j+(T_A-1)} (1-\mu_A)^{(t_k-t_j)} \times \mu_A \times \sum_{y_2=t_k}^{t_k+(T_A-1)} (1-\mu_A)^{(y_2-t_k)} \times \begin{pmatrix} \mathbb{I}[(t_2 - T_E - T_L - T_P - T_A) < y_2 \\ \leq (t_2 - T_E - T_L - T_P)] \\ \times \beta \times (1-\mu_E)^{T_E} \\ \times (1-\mu_L)^{T_L} \times (1-\mu_P)^{T_P} \\ \times (1-\mu_A)^{(t_2-y_2-T_E-T_L-T_P)} \end{pmatrix}.$$

This is the same equation as for the larva-larva case with two exceptions: i) daily egg production is multiplied by the proportion of eggs that survive the egg, larva, pupa and adult stages up to the day of sampling to reflect the fact that adults rather than larvae are being sampled, and ii) the indicator function limits consideration to cases where the day of sibling egg-laying, $y_2$, is between days $(t_2 - T_E - T_L - T_P - T_A)$ and $(t_2 - T_E - T_L - T_P)$, again accounting for the increased time elapsed between egg-laying and adult sampling.

Next, we consider the larva-adult half-sibling kinship probability, $P_{HSLA}(t_2|t_1)$, which represents the probability that, given a larva sampled on day $t_1$, an adult sampled on day $t_2$ is their half-sibling:

$$P_{HSLA}(t_2|t_1) = \frac{\mathbb{E}\left[\text{Adults at time } t_2 \text{ that are half-siblings of a larva sampled at time } t_1\right]}{\mathbb{E}\left[\text{Adult offspring at time } t_2 \text{ from all adult females}\right]} = \frac{E_{HSLA}(t_2|t_1)}{E_A}.$$

Here, $E_{HSLA}(t_2|t_1)$ represents the expected number of surviving adults on day $t_2$ that are half-siblings of a larva sampled on day $t_1$. For convenience, let us refer to the larva sampled on day $t_1$ as individual 1. To calculate $E_{HSLA}(t_2|t_1)$, there are three unknown event times that we treat as latent variables and take an expectation over - i) the day that egg 1 is laid, $y_1$, ii) the day of the mating event between individual 1's mother and father, $t_i$, and iii) the day that individual 1's father emerges as an adult, $t_j$:

$$\begin{aligned} E_{HSLA}(t_2|t_1) = & \sum_{y_1=t_1-T_E-(T_L-1)}^{t_1-T_E} p_L(t_1 - y_1 - T_E) \times \sum_{t_i=y_1-(T_A-1)}^{y_1} p_A(y_1 - t_i) \\ & \times \sum_{t_j=t_i-(T_A-1)}^{t_i} p_A(t_i - t_j) \times E_{HSLA}(t_2|t_1, y_1, t_i, t_j). \end{aligned}$$

This is the same equation as for the larva-larva case with the only exception being the term $E_{HSLA}(t_2|t_1, y_1, t_i, t_j)$, which represents the expected number of surviving adults on day $t_2$ that are half-siblings of larva 1, conditional upon egg 1 being laid on day $y_1$, their mother and father mating on day $t_i$, and their father emerging as an adult on day $t_j$. That term is the same as for the adult-adult case, i.e.:

$$E_{HSLA}(t_2|t_1, y_1, t_i, t_j) = E_{HSAA}(t_2|t_1, y_1, t_i, t_j).$$

Next, we consider the adult-larva half-sibling kinship probability, $P_{HSAL}(t_2|t_2)$, which represents the probability that, given an adult sampled on day $t_1$, a larva sampled on day $t_2$ is their half-sibling:

$$P_{HSAL}(t_2|t_1) = \frac{\mathbb{E}\left[\text{Larvae at time } t_2 \text{ that are half-siblings of an adult sampled at time } t_1\right]}{\mathbb{E}\left[\text{Larval offspring at time } t_2 \text{ from all adult females}\right]} = \frac{E_{HSAL}(t_2|t_1)}{E_L}.$$

Here, $E_{HSAL}(t_2|t_1)$ represents the expected number of surviving larvae on day $t_2$ that are half-siblings of an adult sampled on day $t_1$. For convenience, let us refer to the adult sampled on day $t_1$ as individual 1. To calculate $E_{HSAL}(t_2|t_1)$, there are three unknown event times that we treat as latent variables and take an expectation over - i) the day that egg 1 is laid, $y_1$, ii) the day of the mating event between individual 1's mother and father, $t_i$, and iii) the day that individual 1's father emerges as an adult, $t_j$:

$$E_{HSAL}(t_2|t_1) = \sum_{y_1=t_1-T_E-T_L-T_P-(T_A-1)}^{t_1-T_E-T_L-T_P} p_A(t_1-y_1-T_E-T_L-T_P) \times \sum_{t_i=y_1-(T_A-1)}^{y_1} p_A(y_1-t_i) \times \sum_{t_j=t_i-(T_A-1)}^{t_i} p_A(t_i-t_j) \times E_{HSAL}(t_2|t_1,y_1,t_i,t_j).$$

This is the same equation as for the adult-adult case with the only exception being the term $E_{HSAL}(t_2|t_1,y_1,t_i,t_j)$, which represents the expected number of surviving larvae on day $t_2$ that are half-siblings of adult 1, conditional upon egg 1 being laid on day $y_1$, their mother and father mating on day $t_i$, and their father emerging as an adult on day $t_j$. That term is the same as for the larva-larva case, i.e.:

$$E_{HSAL}(t_2|t_1,y_1,t_i,t_j) = E_{HSLL}(t_2|t_1,y_1,t_i,t_j).$$

For pupa-pupa pairs, the half-sibling kinship probability is denoted by $P_{HSPP}(t_2|t_1)$ and represents the probability that, given a pupa sampled on day $t_1$, a pupa sampled on day $t_2$ is their half-sibling. This can be expressed as:

$$P_{HSPP}(t_2|t_1) = \frac{\mathbb{E}\left[\text{Pupae at time } t_2 \text{ that are half-siblings of a pupa sampled at time } t_1\right]}{\mathbb{E}\left[\text{Pupal offspring at time } t_2 \text{ from all adult females}\right]} = \frac{E_{HSPP}(t_2|t_1)}{E_P}.$$

Here, $E_{HSPP}(t_2|t_1)$ represents the expected number of surviving pupae on day $t_2$ that are half-siblings of a pupa sampled on day $t_1$. For convenience, let us refer to the pupa sampled on day $t_1$ as individual 1. To calculate $E_{HSPP}(t_2|t_1)$, there are three unknown event times that we treat as latent variables and take an expectation over - i) the day that egg 1 is laid, $y_1$, ii) the day of the mating event between individual 1's mother and father, $t_i$, and iii) the day that individual 1's father emerges as an adult, $t_j$:

$$E_{HSPP}(t_2|t_1) = \sum_{y_1=t_1-T_E-T_L-(T_P-1)}^{t_1-T_E-T_L} p_P(t_1-y_1-T_E-T_L) \times \sum_{t_i=y_1-(T_A-1)}^{y_1} p_A(y_1-t_i) \times \sum_{t_j=t_i-(T_A-1)}^{t_i} p_A(t_i-t_j) \times E_{HSPP}(t_2|t_1,y_1,t_i,t_j).$$

This is the same equation as for the larva-larva case with two exceptions: i) the expectation over the day that individual 1 is laid is taken over days $(t_1-T_E-T_L-(T_P-1))$ through $(t_1-T_E-T_L)$ to account for the additional time elapsed between the larva and pupa life stages, and ii) the probability that egg 1 is laid on day $(t_1-y_1-T_E-T_L)$, $p_P(t_1-y_1-T_E-T_L)$, reflects the pupa age probability distribution due to this being the relevant life stage. The term $E_{HSPP}(t_2|t_1,y_1,t_i,t_j)$ represents the expected number of surviving pupae on day $t_2$ that are half-siblings of pupa 1, conditional upon egg 1 being laid on day $y_1$, their mother and father mating on day $t_i$, and their father emerging as an adult on day $t_j$. This is given by:

$$E_{HSPP}(t_2|t_1,y_1,t_i,t_j) = \sum_{t_k=t_j}^{t_j+(T_A-1)} (1-\mu_A)^{(t_k-t_j)} \times \mu_A \times \sum_{y_2=t_k}^{t_k+(T_A-1)} (1-\mu_A)^{(y_2-t_k)} \times \begin{pmatrix} \mathbb{I}[(t_2-T_E-T_L-T_P) < y_2 \\ \leq (t_2-T_E-T_L)] \\ \times \beta \times (1-\mu_E)^{T_E} \times (1-\mu_L)^{T_L} \\ \times (1-\mu_P)^{(t_2-y_2-T_E-T_L)} \end{pmatrix}.$$

This is the same equation as for the larva-larva case with two exceptions: i) daily egg production is multiplied by the proportion of eggs that survive the egg, larva and pupa stages up to the day of sampling to reflect the fact that pupae rather than larvae are being sampled, and ii) the indicator function limits consideration to cases where the day of sibling egg-laying, $y_2$, is between days $(t_2 - T_E - T_L - T_P)$ and $(t_2 - T_E - T_L)$, again accounting for the additional time elapsed between the larva and pupa life stages.

Next, we consider the pupa-larva half-sibling kinship probability, $P_{HSPL}(t_2|t_1)$, which represents the probability that, given a pupa sampled on day $t_1$, a larva sampled on day $t_2$ is their half-sibling:

$$P_{HSPL}(t_2|t_1) = \frac{\mathbb{E}\,[\text{Larvae at time } t_2 \text{ that are half-siblings of a pupa sampled at time } t_1]}{\mathbb{E}\,[\text{Larval offspring at time } t_2 \text{ from all adult females}]} = \frac{E_{HSPL}(t_2|t_1)}{E_L}.$$

Here, $E_{HSPL}(t_2|t_1)$ represents the expected number of surviving larvae on day $t_2$ that are half-siblings of a pupa sampled on day $t_1$. For convenience, let us refer to the pupa sampled on day $t_1$ as individual 1. To calculate $E_{HSPL}(t_2|t_1)$, there are three unknown event times that we treat as latent variables and take an expectation over - i) the day that egg 1 is laid, $y_1$, ii) the day of the mating event between individual 1's mother and father, $t_i$, and iii) the day that individual 1's father emerges as an adult, $t_j$:

$$E_{HSPL}(t_2|t_1) = \sum_{y_1=t_1-T_E-T_L-(T_P-1)}^{t_1-T_E-T_L} p_P(t_1-y_1-T_E-T_L) \times \sum_{t_i=y_1-(T_A-1)}^{y_1} p_A(y_1-t_i) \\ \times \sum_{t_j=t_i-(T_A-1)}^{t_i} p_A(t_i-t_j) \times E_{HSPL}(t_2|t_1,y_1,t_i,t_j).$$

This is the same equation as for the pupa-pupa case with the only exception being the term $E_{HSPL}(t_2|t_1,y_1,t_i,t_j)$, which represents the expected number of surviving larvae on day $t_2$ that are half-siblings of pupa 1, conditional upon egg 1 being laid on day $y_1$, their mother and father mating on day $t_i$, and their father emerging as an adult on day $t_j$. That term is the same as for the larva-larva case, i.e.:

$$E_{HSPL}(t_2|t_1,y_1,t_i,t_j) = E_{HSLL}(t_2|t_1,y_1,t_i,t_j).$$

Next, we consider the larva-pupa half-sibling kinship probability, $P_{HSLP}(t_2|t_1)$, which represents the probability that, given a larva sampled on day $t_1$, a pupa sampled on day $t_2$ is their half-sibling:

$$P_{HSLP}(t_2|t_1) = \frac{\mathbb{E}\,[\text{Pupae at time } t_2 \text{ that are half-siblings of a larva sampled at time } t_1]}{\mathbb{E}\,[\text{Pupal offspring at time } t_2 \text{ from all adult females}]} = \frac{E_{HSLP}(t_2|t_1)}{E_P}.$$

Here, $E_{HSLP}(t_2|t_1)$ represents the expected number of surviving pupae on day $t_2$ that are half-siblings of a larva sampled on day $t_1$. For convenience, let us refer to the larva sampled on day $t_1$ as individual 1. To calculate $E_{HSLP}(t_2|t_1)$, there are three unknown event times that we treat as latent variables and take an expectation over - i) the day that egg 1 is laid, $y_1$, ii) the day of the mating event between individual 1's mother and father, $t_i$, and iii) the day that individual 1's father emerges as an adult, $t_j$:

$$E_{HSLP}(t_2|t_1) = \sum_{y_1=t_1-T_E-(T_L-1)}^{t_1-T_E} p_L(t_1-y_1-T_E) \times \sum_{t_i=y_1-(T_A-1)}^{y_1} p_A(y_1-t_i) \times \sum_{t_j=t_i-(T_A-1)}^{t_i} p_A(t_i-t_j) \times E_{HSLP}(t_2|t_1,y_1,t_i,t_j).$$

This is the same equation as for the larva-larva case with the only exception being the term $E_{HSLP}(t_2|t_1,y_1,t_i,t_j)$, which represents the expected number of surviving pupae on day $t_2$ that are half-siblings of larva 1, conditional upon egg 1 being laid on day $y_1$, their mother and father mating on day $t_i$, and their father emerging as an adult on day $t_j$. That term is the same as for the pupa-pupa case, i.e.:

$$E_{HSLP}(t_2|t_1,y_1,t_i,t_j) = E_{HSPP}(t_2|t_1,y_1,t_i,t_j).$$

Next, we consider the pupa-adult half-sibling kinship probability, $P_{HSPA}(t_2|t_1)$, which represents the probability that, given a pupa sampled on day $t_1$, an adult sampled on day $t_2$ is their half-sibling:

$$P_{HSPA}(t_2|t_1) = \frac{\mathbb{E}\left[\text{Adults at time } t_2 \text{ that are half-siblings of a pupa sampled at time } t_1\right]}{\mathbb{E}\left[\text{Adult offspring at time } t_2 \text{ from all adult females at consistent times}\right]} = \frac{E_{HSPA}(t_2|t_1)}{E_A}.$$

Here, $E_{HSPA}(t_2|t_1)$ represents the expected number of surviving adults on day $t_2$ that are half-siblings of a pupa sampled on day $t_1$. For convenience, let us refer to the pupa sampled on day $t_1$ as individual 1. To calculate $E_{HSPA}(t_2|t_1)$, there are three unknown event times that we treat as latent variables and take an expectation over - i) the day that egg 1 is laid, $y_1$, ii) the day of the mating event between individual 1's mother and father, $t_i$, and iii) the day that individual 1's father emerges as an adult, $t_j$:

$$E_{HSPA}(t_2|t_1) = \sum_{y_1=t_1-T_E-T_L-(T_P-1)}^{t_1-T_E-T_L} p_P(t_1-y_1-T_E-T_L) \times \sum_{t_i=y_1-(T_A-1)}^{y_1} p_A(y_1-t_i) \times \sum_{t_j=t_i-(T_A-1)}^{t_i} p_A(t_i-t_j) \times E_{HSPA}(t_2|t_1,y_1,t_i,t_j).$$

This is the same equation as for the pupa-pupa case with the only exception being the term $E_{HSPA}(t_2|t_1,y_1,t_i,t_j)$, which represents the expected number of surviving adults on day $t_2$ that are half-siblings of pupa 1, conditional upon egg 1 being laid on day $y_1$, their mother and father mating on day $t_i$, and their father emerging as an adult on day $t_j$. That term is the same as for the adult-adult case, i.e.:

$$E_{HSPA}(t_2|t_1,y_1,t_i,t_j) = E_{HSAA}(t_2|t_1,y_1,t_i,t_j).$$

Finally, we consider the adult-pupa half-sibling kinship probability, $P_{HSAP}(t_2|t_1)$, which represents the probability that, given an adult sampled on day $t_1$, a pupa sampled on day $t_2$ is their half-sibling:

$$P_{HSAP}(t_2|t_1) = \frac{\mathbb{E}\left[\text{Pupae at time } t_2 \text{ that are half-siblings of an adult sampled at time } t_1\right]}{\mathbb{E}\left[\text{Pupal offspring at time } t_2 \text{ from all adult females}\right]} = \frac{E_{HSAP}(t_2|t_1)}{E_P}.$$

Here, $E_{HSAP}(t_2|t_1)$ represents the expected number of surviving pupae on day $t_2$ that are half-siblings of an adult sampled on day $t_1$. For convenience, let us refer to the adult sampled on day $t_1$ as individual 1. To calculate $E_{HSAP}(t_2|t_1)$, there are three unknown event times that we treat as latent variables and take an expectation over - i) the day that egg 1 is laid, $y_1$, ii) the day of the mating event between individual 1's mother and father, $t_i$, and iii) the day that individual 1's father emerges as an adult, $t_j$:

$$E_{HSAP}(t_2|t_1) = \sum_{y_1=t_1-T_E-T_L-T_P-(T_A-1)}^{t_1-T_E-T_L-T_P} p_A(t_1 - y_1 - T_E - T_L - T_P) \times \sum_{t_i=y_1-(T_A-1)}^{y_1} p_A(y_1 - t_i) \times \sum_{t_j=t_i-(T_A-1)}^{t_i} p_A(t_i - t_j) \times E_{HSAP}(t_2|t_1, y_1, t_i, t_j).$$

This is the same equation as for the adult-adult case with the only exception being the term $E_{HSAP}(t_2|t_1, y_1, t_i, t_j)$, which represents the expected number of surviving pupae on day $t_2$ that are half-siblings of adult 1, conditional upon egg 1 being laid on day $y_1$, their mother and father mating on day $t_i$, and their father emerging as an adult on day $t_j$. That term is the same as for the pupa-pupa case, i.e.:

$$E_{HSAP}(t_2|t_1, y_1, t_i, t_j) = E_{HSPP}(t_2|t_1, y_1, t_i, t_j).$$
